# Axon morphology is modulated by the local environment and impacts the non-invasive investigation of its structure-function relationship

**DOI:** 10.1101/2020.05.29.118737

**Authors:** Mariam Andersson, Hans Martin Kjer, Jonathan Rafael-Patino, Alexandra Pacureanu, Bente Pakkenberg, Jean-Philippe Thiran, Maurice Ptito, Martin Bech, Anders Bjorholm Dahl, Vedrana Andersen Dahl, Tim B. Dyrby

**Author notes:** **Corresponding authors**, Mariam Andersson & Tim B. Dyrby.

## Abstract

Axonal conduction velocity, which ensures efficient function of the brain network, is related to axon diameter. Non-invasive, in vivo axon diameter estimates can be made with diffusion magnetic resonance imaging, but the technique requires 3D validation. Here, high resolution, 3D synchrotron X-ray Nano-Holotomography images of white matter samples from the corpus callosum of a monkey brain reveal that blood vessels, cells and vacuoles affect axonal diameter and trajectory. Within single axons, we find that the variance in diameter and conduction velocity correlates with the mean diameter, contesting the value of precise diameter determination in larger axons. These complex 3D axon morphologies drive previously reported 2D trends in axon diameter and g-ratio. Furthermore, we find that these morphologies bias the estimates of axon diameter with diffusion magnetic resonance imaging and, ultimately, impact the investigation and formulation of the axon structure-function relationship.

## 1. Introduction

Axons form the communication infrastructure of the brain network. Upon their initial discovery by Otto Friedrich Karl Deiters circa 1860, they were described as “cylinders” – a description still widely used today. The saltatory conduction velocity (CV) of action potentials along myelinated axons depends on their morphology, including axon diameter^1^ (AD) and thickness of the CV-boosting, insulating myelin sheath^2^. The ratio of AD to the outer fiber diameter, the g-ratio, is thought to be fixed to an optimal value that promotes high CVs and minimizes energy consumption^3^. Simulation studies^4,5^ predict optimal g-ratios around 0.7 in the central nervous system (CNS), matching histological data^6,7^. The concept of a given g-ratio along straight, cylindrical axons enables an inference of the outer fiber diameter from the inner AD, allowing classical structure-function relations between outer fiber diameter and CV^8,9^ to be reformulated in terms of AD and a constant g-ratio^8,9^. This makes an investigation of brain network function accessible with techniques that can measure AD.

Histological tracer studies of axons between brain sites reveal that the diameter, and thus CV, of an axon depends on its origin^10^ and target^7^, corroborating the functional significance of AD. AD is potentially also a biomarker for neurodegenerative diseases like Multiple Sclerosis (MS), which has been shown to preferentially attack smaller axons^11^. To provide useful diagnostic information, the white matter (WM) microstructure and axon morphology must be characterized in vivo. Diffusion Magnetic Resonance Imaging (MRI) uses the diffusion of water molecules to non-invasively probe the WM microstructure in the living brain. Although MRI voxels are typically on the scale of ~1 mm, it is possible to estimate axonal dispersion^12^, the axon diameter distribution (ADD)^13^ and the mean of the ADD^14^ by fitting 3D biophysical models to the acquired diffusion signal^15^. However, diffusion MRI-based AD estimates^14,16,17^ are larger than those obtained by histology^15,18^. A potential cause is inaccurate modeling of the WM compartments, including the century old representation of myelinated axons as cylinders. A validation of the 3D WM anatomy could thus improve diffusion MRI-based AD estimations^17^ and shed light on the validity of enforcing a cylindrical geometry and constant g-ratio in axonal structure-function relations. Recent 3D electron microscopy (EM) studies on axon morphology of the mouse reveal, in high resolution, non-uniform ADs and trajectories^19,20^. However, axons are only tracked for up to 20 μm, a fraction of their length in MRI voxels.

Here, we characterize the long-range micro-morphologies of axons against the backdrop of the complex 3D WM environment consisting of blood vessels, cells and vacuoles. With synchrotron X-ray Nano-Holotomography (XNH), we acquire MRI measurements of the WM from the same monkey brain as in Alexander et al. (2010)^14^ and Dyrby et al. (2013)^21^, in which the MRI-derived AD estimates were larger than those estimated by histology. The 3D WM environment is mapped at a voxel size of 75 nm and volume of approximately 150 × 150 × 150 μm^3^. By combining adjacent XNH volumes, we extract axons >660 μm in length and show that AD, axon trajectory and g-ratio depend on the local microstructural environment. The 3D measurements shed light on the interpretation of 2D measurements, highlighting the importance of the third dimension for a robust description of single-axon structure and function. Lastly, by performing Monte Carlo (MC) diffusion simulations on axonal substrates with morphological features deriving from the XNH-segmented axons, we show that geometrical deviations from cylinders cause an overestimation of AD with diffusion MRI.

## 2. Results

### 2.1 Volumetric Mesoscopic White Matter Features

The mean AD indices in the CC were fitted on diffusion MRI volumes of the monkey brain and, as expected, exhibited an increasing trend from the splenium to the genu, but with overestimated diameters^21^ (Figure 1a). In the splenium, along the interhemispheric connection between visual cortices V1/V2, the mean AD index was 1.3 μm. The connection length was delineated by tractography to 49.7 mm (std = 1.9 mm) as described in Methods, resulting in a conduction delay of 4.8 ms.

**Fig 1.**
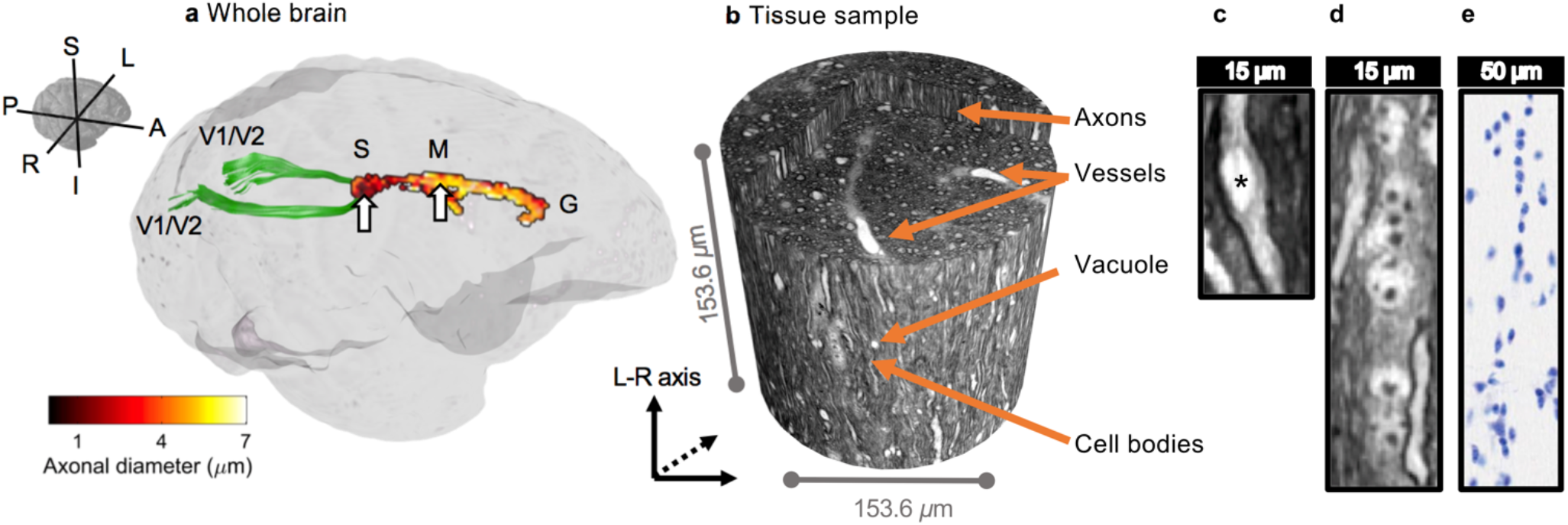
Observed anatomical microstructures within the white matter of the monkey brain. **a** Diffusion-MRI mean AD estimations with the ActiveAx method in the mid-sagittal plane of Corpus callosum (CC) of the monkey brain. Arrows show biopsy locations in the splenium (S) and mid-body (M) regions. The interhemispheric callosal connection between primary visual cortices (V1/V2) is delineated with tractography and is shown in green. **b** 3D XNH volume from the splenium biopsy with an isotropic voxel size of 75 nm, showing detectable anatomical features. The interior of the volume is exposed to reveal the vessels. **c** Close-up of a vacuole (asterisk). **d** Close-up of a cell cluster. **e** Nissl stain light microscopy image showing nuclei in the same splenium region as d in an age matched monkey (BrainMaps: An Interactive Multiresolution Brain Atlas; http://brainmaps.org).

To further investigate the microstructure underlying the diffusion MRI estimates, we extracted and processed cylindrical tissue volumes of 1 mm in diameter from the CC and crossing fiber region for XNH. Four tissue structures were examined, as shown in an XNH volume of the CC splenium in Figures 1b-d: i) long-range myelinated axons ii) blood vessels iii) cells and iv) vacuoles.

*Myelinated axons* were identifiable as bright, tubular shapes with dark contours. The contrast was given by the electron density of the sample, with bright regions corresponding to low density structures, while the dark borders were due to the binding of electron-dense Osmium tetroxide to myelin. Axons exhibited varying diameters throughout the volume. Like axons, *blood vessels* appeared as bright, tubular structures. Their larger diameters and ability to branch distinguished them from axons. Blood cells were rarely detected since they were flushed out during the perfusion process. *Cell nuclei* were distinguishable by their DNA inclusions (Figure 1d). Due to its high electron density, DNA – seen with a Nissl stain in Figure 1e – gave rise to round, dark structures, contained within a less dense nucleoplasm. Generally, the cells clustered and aligned with the axons/blood vessels in the CC samples. *Vacuoles* appeared as hyperintense spheroids and could be located within axons (Figure 1c).

A volumetric quantification of the cell nuclei, vacuoles and blood vessels was performed within an extended cylindrical volume, composed of four stitched XNH volumes having a combined diameter of 153.6 μm and length of 584.5 μm, as in Figure 2a. The tissue compartments were segmented and are shown in Figure 2.

**Fig 2.**
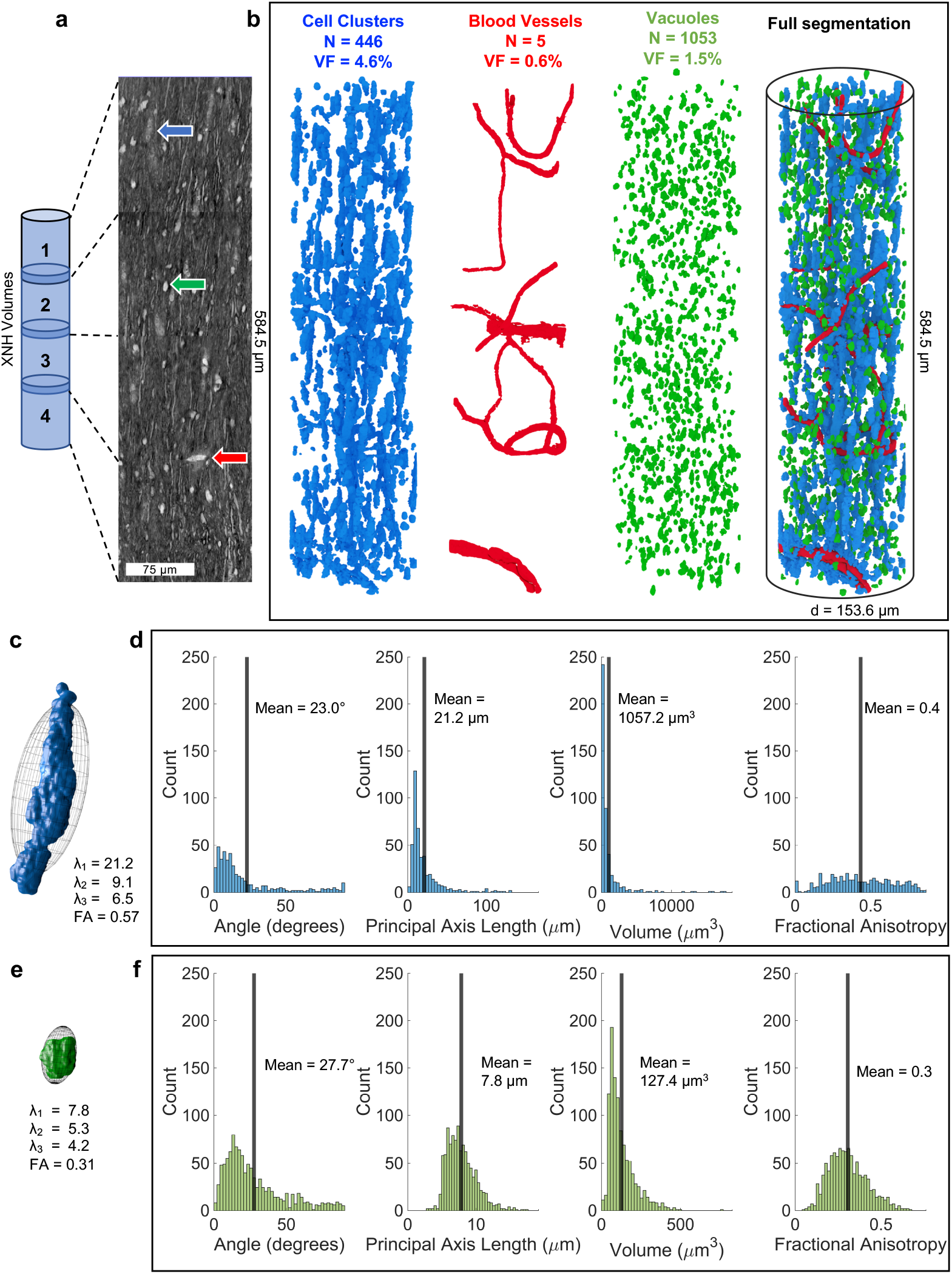
Volumetric quantification of cell clusters, vessels and vacuoles. **a** 2D slice through four stitched XNH volumes (acquired with 13 μm overlap) of the monkey brain splenium. Blue arrow, cell body. Green arrow, vacuole. Red arrow, blood vessel. **b** 3D reconstructions of the cell clusters (blue), blood vessels (red) and vacuoles (green) within a cylindrical volume of length 584.4 μm and diameter 153.6 μm. **c, e**. λ_1_, λ_2_ and λ_3_ denote the average first, second and third principal component lengths of tensors fitted to the cell cluster/vacuole structures. These are visualized by ellipsoids and have a fractional anisotropy denoted by FA. In blue/green: examples of an individual cell cluster/vacuole structure. **d, f** Histograms showing the mean values and distributions of: inclination angle (compared to the axon population direction), principal axis length, mean volume and fractional anisotropy across the cellular (blue) and vacuole (green) components respectively.

Generally, the cell clusters, blood vessels and vacuoles were evenly distributed throughout the volumes, as shown in Figure 2b. This was also true for XNH volumes of the midbody and crossing fiber region (results not shown). The average volume fractions of the quantified structures in the splenium were 4.6%, 0.6% and 1.5% for the cell clusters, blood vessels and vacuoles respectively. The ECS could not be distinguished. Since the samples were dehydrated during tissue processing, it may have shrunk considerably, and the remaining volume fraction is thought to be occupied mostly by myelinated axons. The blood vessels were few, but occupied the largest volume fraction after the axons with diameters between 4 and 10 μm.

Morphological characteristics of the cell clusters are shown in Figure 2d. The cell nuclei had a mean diameter of 5.5 μm (std = 0.73 μm, *N* = 38). Assuming spherical nuclei, the average cell cluster of volume 1057 μm^3^ contained 12 cells. They could be represented by tensors whose principal axes often aligned with the axons (Figure 2d) or nearest blood vessels. The typical cell cluster tensor shape, produced by averaging the first, second and third principal axis lengths of all clusters, is shown in Figure 2c and has a fractional anisotropy (FA) of 0.57 and principal axis lengths between 6.5 and 21.2 μm. The morphological characteristics of the considerably smaller vacuoles are shown in Figure 2f. The typical vacuole tensor shape (Figure 2e) had FA = 0.31 and principal axis lengths ranging between 4.2 and 7.8 μm. The vacuoles were scattered throughout the XNH volume, with some situated within the axons (Figure 1b).

### 2.2 Axonal Micro-Morphology in 3D: Diameters and Dispersion

Axons differed from cylinders in terms of diameter and trajectory changes. Axons (*N=54*) with mean diameters between 2.1 and 3.8 μm were segmented from the XNH volume in Figure 1b as described in Methods. The 54-axon population, shown in Figure 3a, had a mean diameter of 2.7 μm and a volume-weighted mean diameter of 2.9 μm, both significantly larger than the diffusion MRI estimate of 1.3 μm. Smaller axons were observable, but the image resolution and signal-to-noise level of the XNH volumes challenged a robust segmentation. In general, large-diameter (> 2 μm) axons were evenly distributed throughout the tissue at a density of 0.0123/μm^2^, and were surrounded by smaller axons. The mid-body CC and a crossing fiber region were similarly organized (Supplementary Figure 2).

**Fig 3.**
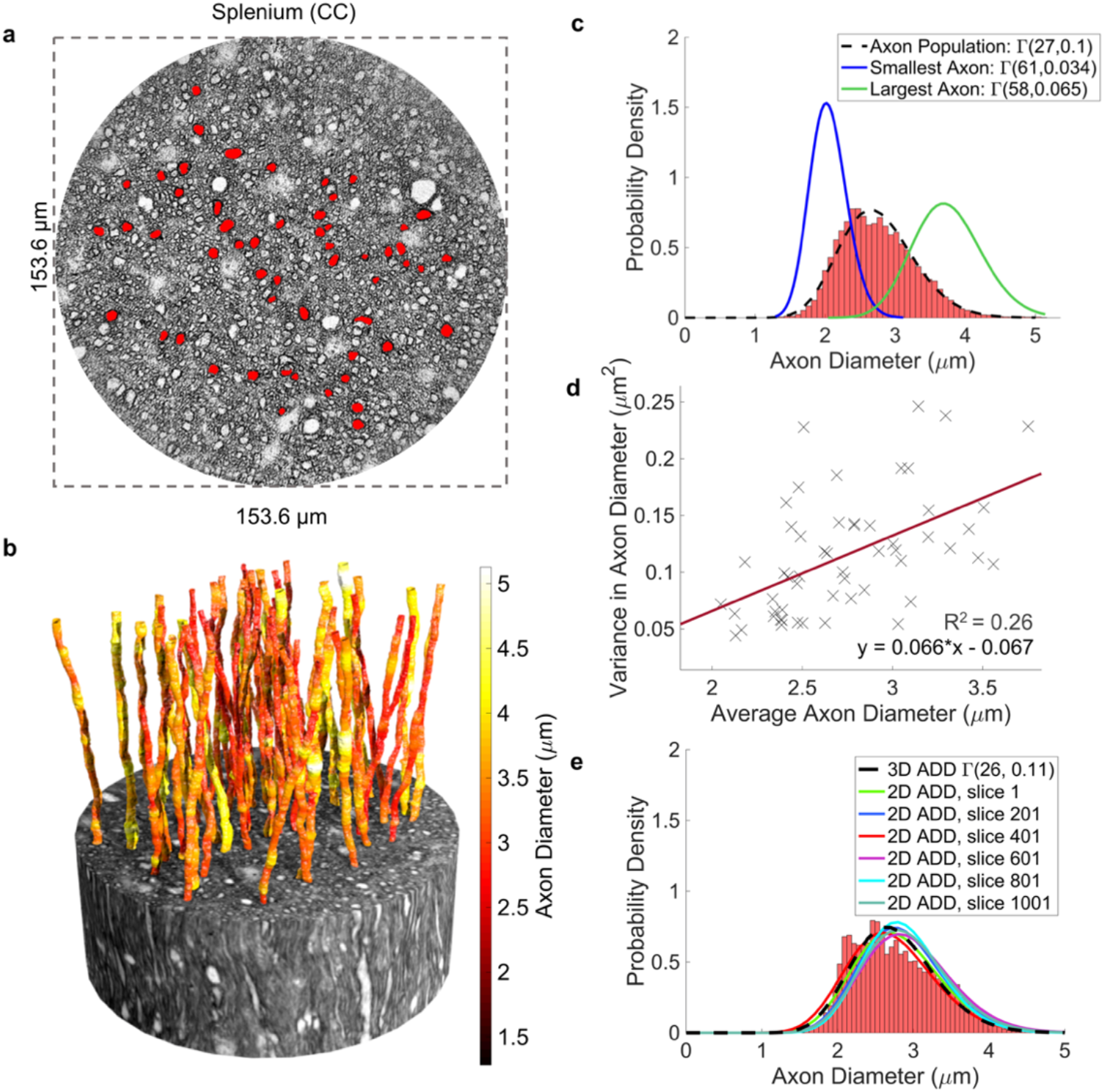
Distribution and morphology of large axons. **a** Distribution of 54 large axons (red) in a 2D slice of the monkey splenium. **b** 54 segmented axons from the monkey splenium. The axon lengths range from 124-170 μm, with average ADs between 2.1 and 3.8 μm. **c** Combined 3D ADD consisting of diameter measurements every 150 nm along all 54 axons and associated Gamma distribution fit (black). For comparison, the fits of the longitudinal ADDs of the thinnest (blue) and thickest axon (red) are shown. **d** The variance in longitudinal axon diameter correlates positively with average axon diameter. **e** Histogram and corresponding Gamma distribution fit (black striped line) of combined 3D ADD, along with Gamma distribution fits of the 2D ADD, sampled every 200 slices of the image sub-volume in which all 54 axons were present.

#### 2.2.1 Longitudinal AD variations

The trajectories of the 54 segmented axons, shown in Figure 3b, ranged in length between 124 and 170 μm, with a combined length of 8.36 mm. Their diameters varied non-systematically along their lengths. We define AD to be the *equivalent diameter*, the diameter of a circle with the same area as the axonal cross section perpendicular to its local trajectory, as in Abdollahzadeh et al. (2019)^19^. The combined 3D ADD, representing all diameters measured at 150 nm intervals along all 54 axons, had a mean diameter of 2.7 μm, and followed a gamma distribution with parameters *a* = 27 and *b* = 0.2 (Figure 3c).

Gamma distributions were also fitted to the individual longitudinal ADDs of the largest and smallest mean diameter-axons respectively. The longitudinal ADD of the largest axon was similar in width to that of the combined 3D ADD, while the smallest axon exhibited a significantly narrower distribution (Figure 3c). A weak linear relationship (R^2^ = 0.26) was found between the mean AD and the variance in longitudinal AD (Figure 3d), suggesting that larger axons experience larger diameter variations than small axons, but with significant variability. The maximum encountered variance of 0.25 μm^2^ (Figure 3d) entails that the majority of diameter fluctuations occur within ±0.5 μm of the mean AD.

To compare our measurements with those from 2D techniques, we calculated the slice-wise ADs for the 1139 slices in which all 54 axons were present. The 2D ADDs from six of these slices are shown in Figure 3e and overlap with the combined 3D ADD. The mean 2D diameter of the fiber population throughout their common volume was 2.9 μm; from slice to slice, this value varied at most ±150 nm.

#### 2.2.2 Axonal dispersion

Axonal dispersion is a measure of disorder of the axonal trajectories. On the bundle level, 54 axons, the orientation dispersion (OD) describes the angular spread of the axons around an axis – the main bundle direction. Although axons in the CC were expected to be aligned and straight, the mean OD was 7 degrees in relation to the main bundle direction (Figures 4a-b).

**Fig 4.**
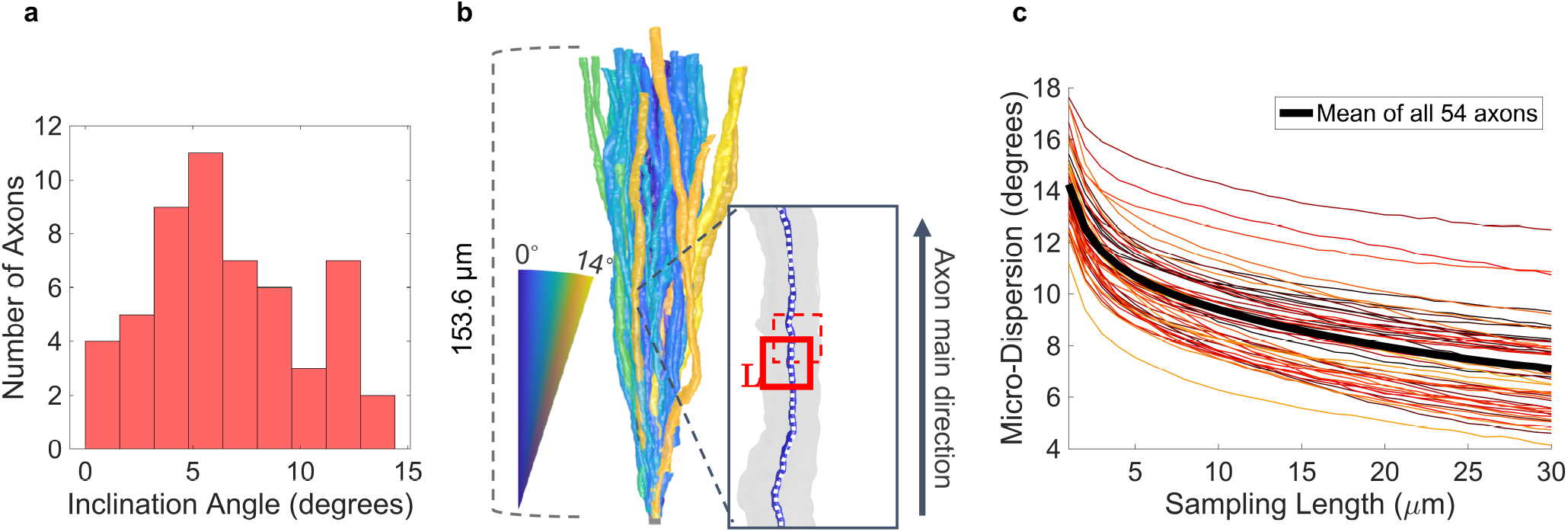
Trajectory variations of large axons. **a** Histogram of the orientation dispersion – the axonal inclination with respect to the main bundle direction – within the axon population. **b** 54 axons from the splenium of the monkey brain translated to a common origin to illustrate the mesoscopic dispersion within the volume. Axon color represents the inclination angle compared to the z-axis. The insert shows the quantification of micro-dispersion along axons: the axon is aligned with the z-axis and a window of length, L, slides along the axon centerline at intervals of L/4. A principal component analysis is performed on points within the window to determine their directionality. The inclination angle to the z-axis is calculated and averaged over all windows. **c** Variation of micro-dispersion relative to main axon direction with sampling length, L.

On the single axon level, we used micro-dispersion to quantify dispersion on different length scales as described in Methods. The micro-dispersion describes the angle between the main axon direction and the direction of axon segments of certain lengths. The micro-dispersion decays smoothly with increasing sampling length (Figure 4c). Between sampling lengths of 1 and 30 μm, the average micro-dispersion decreases from 14 to 7 degrees.

#### 2.2.3 The longitudinal ADD, and not the myelin thickness, dominates longitudinal g-ratio variations

We investigated how the g-ratio and consequently, the CV, varied along axonal internodes – the axon segment between consecutive Nodes of Ranvier – by mapping the long range behavior of six axons (>580 μm length) in the monkey splenium, as shown in Figure 5a. The equivalent inner AD and equivalent outer fiber diameter were evaluated by manual segmentation at *N* randomly generated locations along the axonal internodes (Figure 5b). This revealed a linear correlation between the inner and outer longitudinal diameters, suggesting a constant myelin thickness. The distributions of CVs along the internodes of axons 1, 2 and 6 (Figure 5c) were calculated using the classical relationship^8^ CV = 5.5 · D, where *D* is the outer fiber diameter. Using the tract length of 49.7 mm obtained with tractography, the mean conduction delays along the respective fibers were 2.0, 2.4 and 1.9 ms.

**Fig 5.**
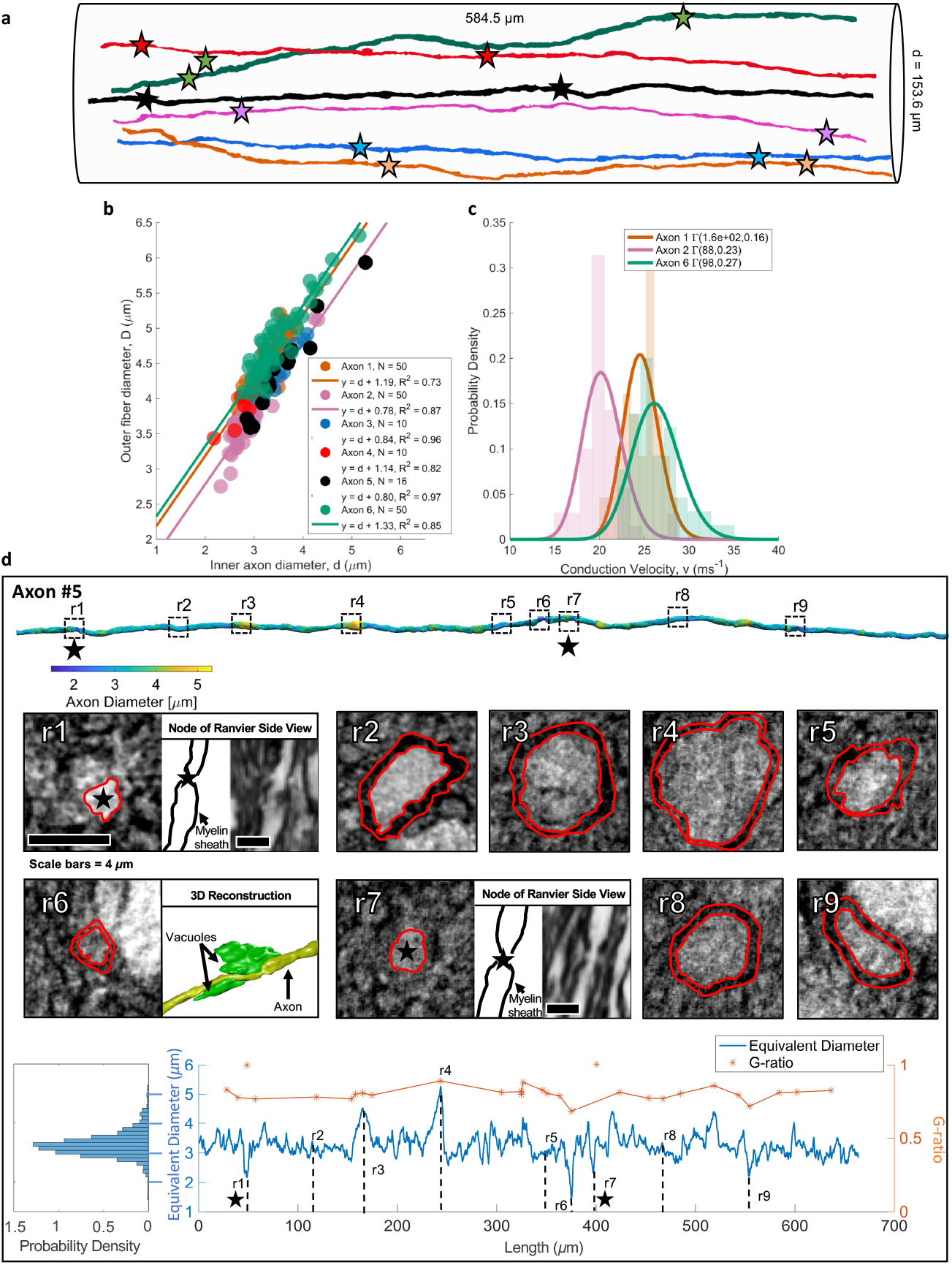
Long-range quantification of axon morphology and myelination. **a** Six long axons segmented from the extended cylindrical XNH volume of the monkey splenium shown in Figure 2a. Different colored stars mark the identified Nodes of Ranvier along each respective axon. **b** Inner AD vs. outer diameter (including thickness of myelin sheath) at *N* randomly sampled points along the internodes of the six axons. Straight lines represent linear fits to the six datasets. The high linear correlation suggests that the myelin thickness remains approximately constant along the internodes, and that the y-intercept is representative of twice the myelin thickness. **c** Histograms and associated gamma distribution fits of CVs along the internodes of Axon 1, 2 and 6. **d** Row 1: segmentation of long axon (number 5, black, in **a** and **b** above), colored according to diameter. Dotted squares marked r1-r9 indicate 9 ROIs. Black stars mark the positions of the Nodes of Ranvier. Row 2: ROI intensity images. The red lines show a manual segmentation of the inner axonal boundary and the outer myelin boundary, respectively. At r1 and r7, there are Nodes of Ranvier which are shown in an orthogonal view. At r6, the axon (in yellow) is squeezed by two vacuoles (in green). Row 3: On left: ADD along axon. Blue line: variation of the longitudinal equivalent diameter, as measured perpendicular to the local trajectory. In orange, the g-ratio for the 9 marked ROIs and 20 additional randomly generated positions along the axon.

Figure 5d depicts the morphological changes occurring over the 662 μm long trajectory of axon 5. Its diameter varied between 1.5 and 5.3 μm, and averaged at 3.3 μm. Of nine selected ROIs labelled r1 to r9, local diameter minima occurred at points r1, r6, r7 and r9. At r1 and r7, we identified Nodes of Ranvier (black stars), separated by 348 μm. At r6, the reduced diameter was caused by two vacuoles. The g-ratio was evaluated at the nine ROIs shown in Figure 5, and at 10 randomly chosen positions inside and outside the internode, respectively. As would be expected of a constant myelin thickness, the g-ratio followed the trend in diameter, with the exception of the Nodes of Ranvier at points r1 and r7.

In contrast to the 54-axon population for which the mean diameter did not deviate more than ±150 nm in any slice of the volume, the mean AD in a single axon could not be reliably established from one measurement. As a measure of stability of the AD, we calculated the cumulative mean AD along axon 5 in Figure 5d. The point at which the cumulative mean diameter became stable to within ±150 nm depended on the position along the axon at which measurements were commenced, and was up to 200 μm for some positions.

### 2.3 Local obstacles such as vacuoles, blood vessels, cell clusters and crossing axon bundles alter the morphology of axons

The presence of certain extra-axonal obstacles gave rise to noticeable trajectory changes. For example, blood vessels visibly warped the surrounding microstructure and disrupted axon trajectories as illustrated in Figures 6a-c.

**Fig 6.**
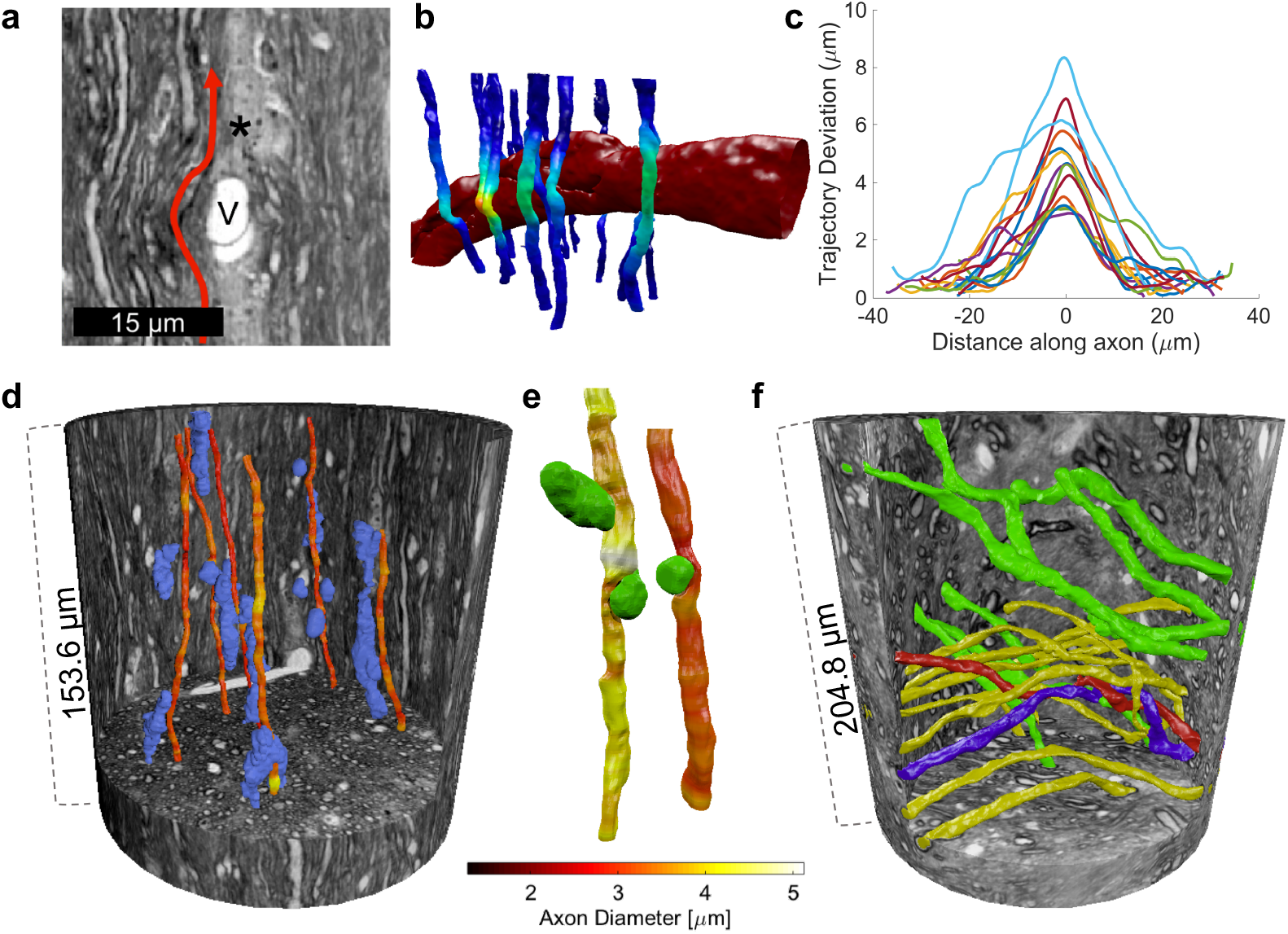
Blood vessels, vacuoles and crossing axons cause axonal trajectory variations. **a** XNH image looking into a vessel, marked V. Cell nuclei, marked by an asterisk, cluster around the vessel. The vessel significantly impacted the nearby axonal trajectories, depicted by the red arrow. **b** 3D reconstruction of **a**. 15 segmented axons are colored according to the deviation from their expected linear trajectories (a linear interpolation of the axon centerline points above and below the blood vessel. Yellow indicates a strong deviation and dark blue indicates little/no deviation. **c** Deviation from expected linear trajectory as a function of distance along the axon, centered on the maximum deviation. **d, e** 3D reconstructions of select cell clusters (blue), vacuoles (green) and axons, whose diameters are given by the colorbar, in the XNH volume of the monkey splenium shown in Figure 3b. The axon trajectories are impacted by the presence of cell clusters, and their diameters and shapes are impacted by neighboring vacuoles. **f** 3D reconstruction of axons in an XNH volume of the crossing fiber region. Two different projection directions are marked by green and yellow. Two axons, colored red and blue, twist around each other.

A sub-volume spanning 690 slices and containing a blood vessel of diameter ~10 μm was isolated (Figure 6a). The expected linear trajectories of the axons, based on their positions in the first and last 75 slices of the volume, were calculated. The axon trajectories exhibited maximum deviations from their expected trajectories between 2 and 9 μm, with the most significant deviations occurring along axon segments within ±10 μm of the blood vessel (Figure 6c). Axonal trajectory changes were also found in response to cell clusters (Figure 6d) and other axons in a crossing fiber region (Figure 6f), in which two axons colored red and blue were found to twist around each other three times within the available volume. Although vacuoles did not cause trajectory variations, they caused a reduction in AD as shown in Figure 5d (point r6) where two vacuoles cause a local decrease in AD, and in Figure 6e.

### 2.4 The influence of axon morphology on diffusion MRI measurements

To assess how axon morphology influences non-invasive diffusion MRI estimates of mean AD, we performed Monte Carlo simulations of the diffusion process in six different 54-axon substrates, see Methods for details. The substrates differed in morphological complexity, ranging from the simplest geometry, G1 – parallel cylinders, to G6 – the XNH segmentation (Figure 7a). Each axon in each substrate inherited its morphological properties (OD, micro-dispersion and mean diameter/longitudinal diameter) from the respective XNH-segmented axons.

**Figure 7.**
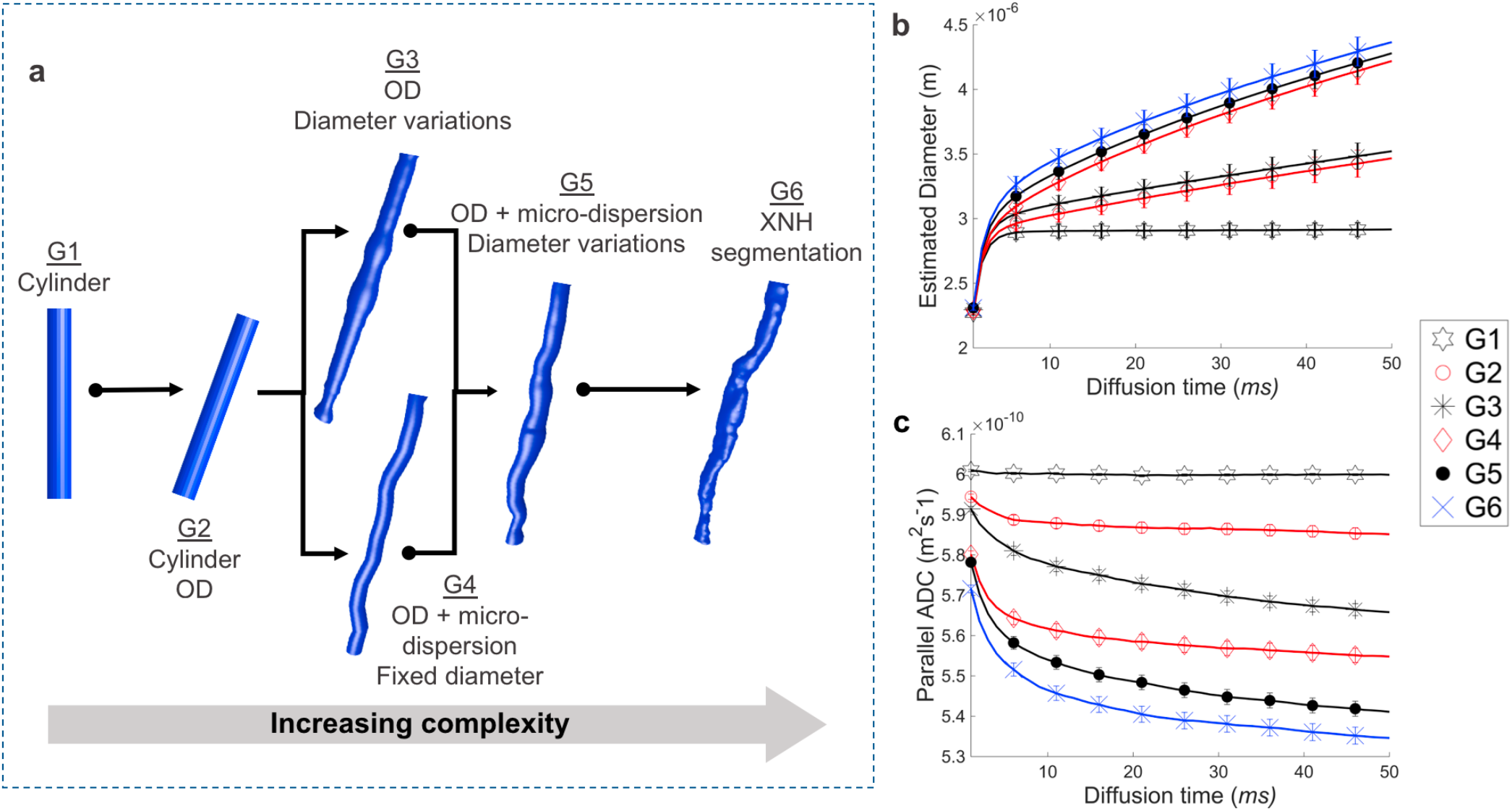
Examining the diffusion properties of different axonal geometries. **a** The morphological features of the XNH-segmented axons in Figures 3 and 4 are directly mapped to the six different axon classes generated for Monte Carlo simulations. G1 – straight cylinders of diameter corresponding to XNH axon mean diameters, G2 – same as G1 plus segmented OD, G3 – segmented OD and longitudinal ADD, G4 – segmented OD and micro-dispersion, G5 – segmented OD, micro-dispersion and longitudinal ADD, and G6, the XNH segmentation. **b** The variation of estimated AD and, **c,** parallel intra-axonal ADC with diffusion time for geometries G1-G6. Error bars represent the standard error, reflecting the spread in diameters/ADCs across the individual axons.

After an initial diffusion time of 6 ms, the mean AD in the parallel cylinders remained constant, as expected, at the true volume-weighted mean AD of 2.9 μm. For the other substrates, however, AD was overestimated. The degree of overestimation increased with diffusion time and substrate complexity (Figure 7b). The simulations also demonstrated the effects of OD (G2) on AD, and revealed that the overestimation of AD is more significant when there is micro-dispersion than diameter variations (G3 vs G4).

The parallel Apparent Diffusion Coefficient (ADC) describes the rate of diffusion along a cylinder, or axon, and is often assumed to be a fixed value in MRI-based AD estimation. Our results show parallel ADC in parallel cylinders (G1) was constant and equal to the chosen intrinsic diffusivity of 6·10^−10^ m^2^/s. However, for substrates that exhibited OD, micro-dispersion and/or diameter variations, the parallel ADC decreased with diffusion time and substrate complexity. Substrates with longitudinal AD variations (G4, G5, G6) exhibited particularly steep decreases in the parallel ADC with diffusion time. The 3D effects of AD variation and dispersion (OD and micro-dispersion) could thus be distinguished from each other based on the time-dependence of the parallel ADC.

## 3. Discussion

By performing high resolution 3D X-ray nano-holotomography on intact white matter samples from a monkey brain, we demonstrate for the first time the interplay between extra-axonal structures (blood vessels, cells, vacuoles) and the micro-morphology of axons. Contrary to Deiters’ century-old description of axons as cylinders, we find that axon diameter and trajectory vary along the length of the axon, often due to obstacles in the local microstructural environment. These morphological changes entail that large axons are non-specific in terms of diameter and conduction velocity. We thus question the value of precisely measuring their diameters and the validity of enforcing cylindrical geometries in axonal structure-function relationships. Further, we show that the 3D morphologies of axons may drive previously reported trends in 2D axon diameter distributions and g-ratio distributions. Our results have significant impact for axon diameter determination with 2D techniques and – as we show here – diffusion MRI. We foresee that a thorough morphological characterization of axons and their structural context will guide the non-invasive investigation of axon morphology with diffusion MRI and, consequently, the investigation of brain network function.

### 3.1 The non-specificity of the diameters and g-ratios of large axons

Single axons lack well defined diameters, and this has implications for the interpretation of 2D ADDs. Our findings show that individual axons exhibit longitudinal ADDs whose widths correlate with AD. Controversially, single axons are not always well described by classical 2D histological measurements. For one 662 μm long axon, we show that a robust characterization of its mean diameter demands that AD be sampled for up to 200 μm at intervals of 0.15 μm. However, we find the opposite for a quantification of population mean AD. In axon populations, the 2D slice-wise ADDs match the 3D ADD, indicating that the 3D longitudinal AD variations are represented in the 2D ADDs. This has two consequences. Firstly, it entails that previously reported 2D ADDs from EM^22^ and light microscopy^7^ also reflect the 3D ADD of the axon population if quantified over a sufficiently large volume. Secondly, contrary to the traditional interpretation of the ADD, it implies that the 2D ADDs are not directly representative of individual ADs. Instead, the 2D ADD may be interpreted as the *sampling* of the longitudinal ADD of each axon. We thus propose that the characteristic tail of the 2D ADD may arise as a result of the broad longitudinal ADDs of large axons. Therefore, observations of “giant axons”^15,22^ may not indicate the presence of very large axons, but instead the local AD variations along large axons.

The morphological non-specificity of large axons may indicate that they have similar functions, and that it is not important to precisely determine their diameters. The wide and overlapping longitudinal ADDs of large axons entail that their diameters are less defined than those of smaller axons, which exhibit narrower longitudinal ADDs. Given these effects, we suggest a categorization of axons into “large” and “small”. Similar categorizations are made in the peripheral nervous system (PNS). The fast-conducting, larger Alpha/Beta axons of the sciatic nerve have a broad ADD which partially overlaps the narrower ADD of the smaller, slower C and Alpha/Delta fibers^23^. Although the structure-function relationship of CNS axons is not as well described as that of the sciatic nerve, it has been shown that CNS AD correlates with the neuron soma size^18,24^. ADs may thus be cell-type specific. The possibility that smaller and larger axons may encode different functional features^25^ further supports a size-specific axon categorization. Hence, we suggest that it may be more important to perform a size-specific axon categorization than to measure precise mean ADs, as done today with histology and diffusion MRI techniques.

The longitudinal ADDs drive variations in g-ratio, while myelin remains stable between Nodes of Ranvier. Along *single* internodes, we found a linear relationship between the inner AD and outer fiber diameter, entailing a non-linear relationship between g-ratio and AD. Interestingly, this trend is of similar shape to that observed in 2D EM measurements of g-ratios in *populations* of CNS axons^26^. For such 2D measurements, Berthold et al (1983) formulated a log-linear relationship between the number of myelin lamellae (proportional to myelin thickness, assuming a constant lamellar spacing) and the inner AD^27^. For single axons, however, we did not find that the myelin thickness varied as a function of local AD within internodes. Instead, the linear correlation between the longitudinal inner AD and outer fiber diameter indicated that the myelin thickness along internodes was constant. Deviations from the linear trend could be due to too few sampling points, the myelin wrapping mechanism^28^, measurement uncertainty in the inner and outer ADs, or myelin-disturbing vacuoles or fixation effects. Histological studies suggest that the myelin thickness shows a stronger correlation to CV than the AD^2^, while modelling studies report that myelin is expensive to remodel^29^. A constant myelin thickness along single internodes is thus motivated from a signal timing and energy perspective, entailing that the longitudinal g-ratio variation can be driven by the longitudinal ADD. We expect the 2D g-ratio distributions to represent a sampling of the 3D longitudinal g-ratio distribution, in the same way as the 2D ADDs reflect the longitudinal AD variations. Consequently, Berthold’s log-linear relationship relating the number of myelin lamellae to the AD may also be partially driven by longitudinal variations in AD. Furthermore, although large g-ratios can be detected in 2D images, these may be confined to local axonal segments. G-ratio measurements of individual axons are therefore not meaningful on their own, since they capture only a snapshot of the 3D axon morphology. Indeed, our findings here apply only to large axons with diameters >2 μm, and smaller axons remain to be investigated.

Longitudinally varying axon morphologies suggest that individual, myelinated axons exhibit varying CVs. The calculated mean conduction delays along three axons in the interhemispheric, callosal V1/V2 connection were between 1.9 to 2.4 ms, contrasting with our diffusion MRI-estimated conduction delay of 4.8 ms. The longer delay of 4.8 ms agrees with the calculated conduction delay within the same tract in the larger macaque brain, where the tract length was estimated with tracers to 63.4 mm, the average AD to be 0.95 μm and the calculated conduction delay to be 4.6 ms (std = 1.2 ms)^30^. We thus conclude that the same tract can contain many different sized axons, resulting in a range of conduction delays. Further demonstrating the nonspecificity of large axons, the range of CVs along single internodes is large, ranging, for example, between 22 to 35 m/s in axon 6 of Figure 5. Additionally, we note that diffusion MRI and tracer-based histology calculations of CV reformulate Waxman’s classical relation to incorporate a constant g-ratio^9,18^, which we show is not the case. Other factors, such as the internodal spacing, also affect the saltatory CV along an entire axon^31^. Nodes of Ranvier were identifiable and internodal distances could be measured in the long axons spanning the four stitched XNH volumes in Figure 2, but such an analysis was beyond the scope of this study. Several studies have investigated the relationship between axon morphology (e.g. AD, node of Ranvier length, internodal length, g-ratio) and CV^3,5,29,32^, but none have yet considered the effects of a longitudinal distribution of g-ratios.

### 3.2 Changes to axonal morphology may be caused by extra-axonal obstacles

The longitudinal variations in AD and trajectory in the vicinity of extra-axonal structures could contribute to the high intracellular volume fraction of around 80% in the brain^33^. In the extended volume of the monkey splenium, the blood vessels, cell clusters and vacuoles accounted for 6.7% of the volume. Aside from the cells clustering parallel to the axons and blood vessels in the CC, the extra-axonal structures occupied space independently of the surrounding axons in the mid-body CC, splenium and crossing fiber region. Morphological variations and spatial distributions of the extra-axonal compartments could not be linked to the presence of axons. On the other hand, the diameter and trajectory variations of axons could be linked to the presence of extra-axonal structures.

Local AD minima were at times associated with the presence of vacuoles. Vacuoles have been reported in EM studies of the healthy CC^19^ and described in^34^. It is possible that vacuolation occurs as a fixation or embedding artifact^35–37^, but we have also observed vacuoles in the unfixed optic nerve of the mouse, studied with cryo-EM (not shown). Reduced ADs were also observed at Nodes of Ranvier, as described in the literature. It has been described that AD is regulated by the axonal cytoskeleton^38^, which maintains axon shape and consists of nanometer thick neurofilaments. Decreases in AD have been shown to occur when axons are subjected to axial or circumferential tension, or microtubule disruption^39^. In the same way, a reduction of tension through disruption of the axonal actin filaments or myosin II causes a diameter increase^39,40^. Adjacent structures such as vacuoles and neighboring axons may place local circumferential tension on the axons, affecting the organization of microtubules and possibly causing the diameter decreases we observe. This could not be confirmed in the XNH volumes as the microtubules and neurofilaments were too small to be resolved. Furthermore, we demonstrate that axonal trajectory changes are caused by blood vessels (Figures 6a-c), cells (Figure 6d) and other crossing axons (Figure 6f). Although axons may have pre-defined targets during axonal growth, we postulate that their trajectories are modified by the extra-axonal environment.

The 54-axon population exhibited both low- and high-frequency trajectory changes. We quantified the dispersion effects across length scales from the microscopic (1 μm) to the mesoscopic (30 μm). Non-straight trajectories are commonly observed in tracer-labelled axonal projections over long distances^9,10,18^, and axonal growth has been shown to follow non-straight trajectories in axons from the frog and chick^17,41^. In our data, the axons skirt around extra-axonal obstacles i.e. cell clusters and blood vessels ranging in size from 4-10 um, and this effect may be a major contributor to the meso-dispersion. The micro-dispersion could be a characteristic of axonal growth, or simply a consequence of the meso-dispersion which, by definition, gives rise to micro-dispersion.

The quantification of axon OD is based on the trajectories of the 54 axons from the CC splenium. These are relatively aligned, and the results from their analysis are supported by inspection of XNH volumes of the CC mid-body. Axons in crossing fiber regions exhibited similar non-straight trajectories and intertwining; such effects have previously been observed histologically in tracer labelled axons^42^. Previously, we have detected indications of local undulations in the CC (results presented in Dyrby et. al, ISMRM 2014, Abstract #2619) with histology, and other studies have derived the theoretical impact of sinusoidal undulation on the diffusion signal for AD estimation^43^. The axon trajectories observed here, however, were irregular. Our 3D XNH data suggests that mesoscopic dispersion can occur due to axons projecting around large obstacles e.g. blood vessels. If these obstacles are aligned and evenly distributed, they may result in the appearance of axonal “undulation”.

### 3.3 Impacts on AD estimation with diffusion MRI

Our findings on the longitudinal ADD and g-ratio variations, as well as the relationship between axon morphology and extra-axonal obstacles, impact microstructural MRI techniques for non-invasive AD estimation. Our MC simulations show that any morphological variation from parallel cylinders incurs an overestimation of AD. This will be the case for all diffusion MRI models that assume a cylindrical axonal geometry and base the AD estimation on measurements perpendicular to the axon population, confirming the need to account for fiber dispersion effects^14,16,21,44^. Some studies implement biophysical models to account for axon dispersion or crossing axon effects^12,45^, but the intra-axonal MRI signal profile intermingles with that of the ECS, challenging a robust fitting^46^. We could not extract the ECS from our XNH data due to insufficient resolution and shrinkage caused by the tissue processing. As such, we simulated only diffusion within the intra-axonal space. Further challenging accurate AD estimates with diffusion MRI, we found that the parallel ADC is time dependent for realistic axonal substrates. Consequently, measured values of the parallel ADC are likely lower than the true intrinsic diffusivity of the tissue. This potentially influences the ActiveAx-estimated mean AD of 1.3 μm which we present here, since the model assumes that the parallel ADC and intrinsic diffusivity of the tissue are equivalent^14^. Simulated substrates with longitudinal AD variations (G3, G5 in Figure 7) exhibited stronger time-dependence in the parallel ADC than substrates with constant ADs (G2, G4), suggesting that diameter variations could be a potential source of the axial time dependence observed in other studies^47^, in line with the suggestions of Fieremans et al. (2016)^48^.

Interestingly, to ensure robust fitting of the axon model to the diffusion MRI data in Alexander et al.(2010) and Dyrby et al. (2013), one had to account for a small “dot” compartment which represented a small fraction of isotropically restricted water molecules. We identified two such possible. These were the vacuoles (1.4% volume fraction) and cell clusters (4.6% volume fraction).

### 3.4 Future directions

Synchrotron XNH of the monkey WM provides anatomical information within volumes approaching the size of MRI voxels, including the micro-morphologies of axons and the volume fractions and morphologies of cells, vacuoles and blood vessels. However, the ECS is not visible, perhaps due to tissue shrinkage, insufficient signal to noise ratio (SNR) and limited resolution at 75 nm. Although scaling factors can be employed to compensate for shrinkage, it is not known if shrinkage affects all WM compartments equally^17^. To fully reconstruct the volume fractions of the respective WM compartments, including the ECS, cryo-techniques could be implemented to preserve the hydrated microstructure. Both cryo-EM^49^ and cryo-XNH provide alternatives to do so.

As with many imaging methods, XNH is subject to a trade-off between the size of the image volume and the resolution. The image quality depends on the imaging time and sample processing, including the sample size, staining, and smoothness of the embedding medium. We chose to image at voxel sizes of 75 nm (splenium) and 100 nm (crossing fiber and mid-body), but the SNR challenged the full segmentation of axons with mean diameters smaller than ~2 μm. The morphological behavior of smaller axons thus remains to be studied. This is possible with XNH, given that one prioritizes resolution and optimizes the sample preparation. However, EM provides superior resolution and there are ongoing efforts to develop large-scale 3D mapping of neural tissue by stacking 2D EM images^50^. Still, EM is time demanding and there is a need to combine techniques that bridge different resolutions and volumes; it would be valuable to image a sample by synchrotron XNH, and thereafter image a sub-region of the same sample with 3D EM.

Lastly, the axonal trajectory variations and dispersion behavior presented here could act as an axonal “fingerprint” to guide the construction of anatomically informed axonal phantoms for MC simulations. Existing frameworks have been developed to model morphological features such as fiber undulation^43,51^ (although we do not observe periodic undulations in our data), and diameter variations^52,53^. Others allow for the generation of a more complex WM environment with beaded axons and cells^54^. None, however, have so far imposed anatomically realistic trajectory patterns or longitudinal ADDs.

## Acknowledgments

M.A. and H.M.K. were supported by the Capital Region Research Foundation (grant number: A5657) (PI:T.D.). M.B. was supported by the Swedish Research Council (grant number E0605401 and E0605402). The authors wish to thank Susanne Sorensen for her assistance with the tissue preparation.

## Author Contributions

Conceptualization, T.B.D., M.A., H.M.K.; Methodology, M.A., H.M.K, J.R.P, A.P, T.B.D., M.B., B.P.; Software, M.A., H.M.K, J.R.P., A.B.D., V.A.D. J.P.T, T.B.D.; Formal Analysis, M.A., H.M.K, J.R.P., T.B.D.; Investigation, M.A., H.M.K, J.R.P, A.P., M.P., A.B.D., V.A.D., T.B.D.; Writing – Original Draft, M.A., T.B.D.; Writing – Review & Editing; M.A., T.B.D, H.M.K., J.R.P., A.P., M.P., B.P., J.P.T., M.B., A.B.D., V.A.D.

## 4. Methods

### 4.1 Monkey brain tissue

The tissue studied came from a 32-month old female perfusion fixated vervet (*Chlorocebus aethiops*) monkey brain, obtained from the Montreal Monkey Brain Bank. The monkey, cared for on the island of St. Kitts, had been treated in line with a protocol approved by The Caribbean Primate Center of St. Kitts. The brain had previously been prepared according to Dyrby et al. (2011)^55^ and ex vivo MRI scanned using the optimized ActiveAx framework for volume-weighted mean AD estimation in Alexander et al.(2010) and Dyrby et al. (2013)^14,21^. Prior to synchrotron X-ray nano-holotomography, the tissue was prepared for whole-brain ex vivo MRI for the estimation of AD and the segmentation of interhemispheric callosal fibers with tractography, as described in the following.

### 4.2 MRI scanning, axon diameter estimation and tractography

#### Tissue preparation for MRI

The fixated tissue was prepared for scanning as described elsewhere^17,55,56^. In short, the tissue was sealed in a double plastic bag with minimal free fluid and placed for temperature stabilization at room temperature overnight prior to scanning. The brain was then positioned in a quadrature volume RF coil on a mechanical stable set up using LEGO^™^ to minimize short-term instabilities, seen as a non-linear decreasing motion artifacts in the first hours of an MRI acquisition^55^. A temperature-stabilized environment was ensured by an constant influx of airflow at room temperature, producing stable diffusion MRI images.

#### MRI scanning

The diffusion MRI dataset was collected on an experimental 4.7 Tesla Agilent MRI scanner with a maximum gradient strength of 600 mT/m. An optimized ActiveAx MRI protocol based on a maximal gradient strength of 300 mT/m for ex vivo tissue as in Dyrby et al. (2013)^21^ was used. The protocol was based on a 2D T2-weighted Pulsed-Gradient-Spin-Echo (PGSE) sequence with a single-line readout to reduce geometric image distortions. The optimized ActiveAx diffusion MRI protocol included three unique b-values [2011, 2957, 9259] s/mm^2^ obtained by the following sequence parameters: Gradient pulse duration (δ) [5.6, 7.0, 10.5] ms, time between onset of gradient pulses (Δ) [12.1, 20.4, 16.9] ms and gradient strength [300, 219, 300] mT/m. The b-values were acquired as shells including [84, 87, 68] non-collinear diffusion weighting encoding directions obtained from the Camino Toolbox^57^. For each shell [15, 16, 13] repeats of b = 0 s/mm^2^ were collected. All shells used the same echo time (TE) of 36 ms and a repetition time (TR) of [4200, 4200, 7200] ms. For whole-brain coverage at an isotropic 0.5 mm^3^ image resolution, a matrix size of 128×256 was used and 100 sagittal slices were acquired in an interleaved manner. The total scan time, excluding a dummy prescan of >6 hours to reduce shortterm instability artifacts, was 50 hours. Prior to axon diameter fitting and tractography the diffusion MRI data sets were denoised^58^ and processed to remove Gibbs ringing artifacts^59^ using the method implementations available in the MRTrix3 software toolbox (RRID: SCR_006971).

#### Axon Diameter estimation

Axon diameter estimation was performed using the ActiveAx framework which is based on the minimal model of white matter diffusion (MMWMD) – a four compartment biophysical model – as described in detail in^14^. In short, the compartments and their respective volume fractions (VF) are:

i. the VF of the extra axonal space modelled as a symmetric tensor.
ii. the VF of intra axonal space modelled as a single cylinder with a mean diameter that best fits the diffusion signal.
iii. the VF of the “dot” compartment reflecting a small fraction of stationary water likely trapped inside vacuoles and cell bodies.
iv. the VF of free diffusion modelled as an isotropic tensor coming from surrounding fluid around the CC.

The MMWMD model was fitted to the diffusion MRI data with the same model parameter settings and multi-stage fitting method as in Dyrby et al. (2013)^21^ using the ActiveAx implementation as part of the Camino Toolbox. Since MRI probes volumes, larger axons contribute more to the intra-axonal signal than smaller ones. Hence, the estimated mean AD using diffusion MRI is a volume-weighted index^14^ A region-of-interest (ROI) was manually drawn to cover the midsagittal plane of the CC within which the volume-weighted mean AD was calculated voxel-wise as the average of 100 repeated estimations.

#### Segmentation of interhemispheric brain connections with tractography-based MRI

Streamline-based tractography was used to segment and estimate the trajectory length of the interhemispheric connection that, via the splenium in the CC, connects the two primary visual cortical areas at the V1/V2 border that represents the vertical in meridian in most species including cats, monkeys and humans^60^. We used a probabilistic streamline approach based on a voxel multi-fiber reconstruction method. First, a constrained spherical deconvolution (CSD) method^61^ implemented in MRTrix3 was fitted to the b=2957 s/mm^2^ shell dataset to give a voxelwise multi-fiber reconstruction of the whole brain. Thereafter, probabilistic streamline tractography was implemented with the SD-STREAM function in MRtrix3 using standard parameters to extract 2000 streamlines^62^. The probabilistic tractography was initialized in a broad seed region covering the splenium region of the midsagittal CC. To obtain only streamlines that projected towards both V1/V2 of each hemisphere, we defined two inclusive regions through which valid streamlines had to project. These regions were manually drawn to cover a broad region in the most anterior coronal slice at which V2 starts. The two inclusive regions were symmetrically in both hemispheres. The seed and inclusive regions were manually defined on the b=0 s mm^−2^ using the FSLeyes software. The mean streamline length and its variation were calculated using the TCKSTATS method in MRTrix3. Visualization of tractography results used MRtrix3 tools and Blender (RRID: SCR_008606).

#### Tractography-based estimation of conduction velocity

To estimate the end-to-end conduction delay of the interhemispheric connection between the V1/V2 border regions and passing through the splenium, the tract was assumed to have a g-ratio, g, of 0.7 as is in other studies^10,30^. The CV was calculated according to^9^ *CV* = (5.5/*g*) · *d*, where *d* is the inner axon diameter. The inner axon diameter was defined to be the axon diameter index obtained by fitting ActiveAx to the diffusion MRI dataset of the monkey brain, as previously described. Together with the tract length from tractography, *L*, the conduction delay (*t*) was given by: *t* = *L/CV*.

### 4.3 Synchrotron XNH imaging and segmentation

#### Tissue preparation for XNH imaging

After MRI acquisition, the whole monkey brain was embedded in agar for mechanical stability during slicing of the brain when placed in a mold. The brain was cut into sagittal slices at thicknesses between 2 and 4 mm. Samples from the midsagittal CC and crossing fiber regions were extracted with a biopsy punch of diameter 1 mm and fixed in 2.5% Glutaraldehyde before staining with 0.5% osmium tetroxide (OsO4) at room temperature for two hours. The stained tissue was dehydrated with an alcohol series and embedded in EPON resin. Excess EPON was removed to produce blocks of approximate dimension 1 x 1 x 4 mm. It was later discovered that the Osmium had not penetrated the tissue sample fully, leaving the core unstained. However, the stained peripheries of the samples were large enough to cover the XNH field of view.

#### S*ynchrotron XNH imaging*

X-ray nano-holotomography (XNH) was performed at beamline ID16A of the European Synchrotron Research Facility (ESRF). Samples were imaged using a nano-focused cone beam^63,64^ of energy 17 keV. Holograms of the samples were recorded at different distances with respect to the focus and the detector to obtain phase maps^64,65^. In practice, sequential tomographic scans were acquired at four different propagation distances by rotating the samples over 180 degrees, and corresponding angular holographic projections were aligned and combined to generate phase maps of the sample. For each tomographic scan, 1800 projections were acquired with exposure times of 0.22 s using a pixel size of 75 nm. The reconstructed image volumes were cylindrical, with height and diameter of 2048 x 2048 x 2048 voxels. The acquisition of one full scan took approximately four hours in total.

#### Segmentation of cell clusters, blood vessels and vacuoles

The intensity distributions of the cell nuclei, blood vessels and vacuoles were wide and overlapping, and the uneven staining complicated a solely intensity-based segmentation. As such, we implemented an in-house intensity- and morphology-based approach in MATLAB to segment the cell nuclei, blood vessels and vacuoles from the four consecutively acquired XNH volumes (Figure 2a).

The segmentation method is based on classic low-level image analysis operations, in the form of intensity thresholds, morphological operations and connected components analysis. Where many conventional methods would attempt to assign labels to all voxels simultaneously or in iterations, we instead solve one class at a time, in the order of their perceived difficulty, starting with the easiest class. The information gained in previous steps serves as fixed prior information in the more difficult later steps e.g., a voxel cannot belong to a particular class, if it has previously been labelled as something else. This approach is taken since it is practically simpler to tune parameters for one class at the time, inspect and manually correct the result until it is satisfactory and then proceed to the next class. Tuning of the parameters and a re-segmentation of the individual volumes could be performed and visualized in a matter of minutes for each compartment. This made it simpler to a) fine-tune the parameters and b) quality control the final solution, which helps to minimize errors.

In order of decreasing average brightness the segmented compartments can be ordered as blood vessel, vacuoles, and cell nuclei. The shape criteria for the different classes were as follows:

- Blood vessels: large, tubular connected regions (volumes > 730 μm^3^), with no specific structures or branching patterns.
- Vacuoles: relatively spherical (minimum elongation, described below, of e = 0.4) with volumes ranging between [15, 600] μm^3^ and spanning maximum 25 μm in the longest direction.
- Cell nuclei clusters: typically appear in relatively large connected regions (estimated to span at least 60 μm in the longest direction with volumes ranging between [75, 2300] μm^3^) and containing dark DNA inclusions.

Suppose that a connected component spans S_CC_ = [s_1_, s_2_, s_3_] unique rows, columns and slices of the volume respectively. The elongation of the component is here calculated as e = inf(S_CC_)/sup(S_CC_). A score of 1 can then roughly be interpreted as a perfect spherical component.

The blood vessels, vacuoles and cell nuclei clusters were large compared to the pixel size of 75 nm. Each of the four XNH volumes was therefore downsampled by averaging by a factor of 5 to improve signal-to-noise ratio and speed up computations. They were then individually segmented to avoid errors due to slightly differing contrasts in the separate volumes. Finally, the segmentations were combined as in Figure 2c, manually corrected in some regions and analyzed by a connected components analysis which rejected the smallest, falsely segmented components.

#### Segmentation of Axon Diameters

Given their varying intensities, morphologies and occasional nodes of Ranvier, segmentation of the axons required a geometrically constrained method, i.e. a segmentation approach which can enforce a closed tubular output. Axons were first preliminarily segmented from downsampled volumes, and thereafter segmented from the high-resolution volumes with voxel size 75 nm.

The image volumes from the XNH were downsampled by averaging by a factor of 5 to give an isotropic voxel size of 375 nm. Each slice was filtered with a Gaussian kernel of width 5 pixels in MATLAB (ver. 2019b). A rough segmentation of the axons was then performed using the adaptive paintbrush in the program ITK-Snap (RRID: SCR_002010). This allowed for the extraction of a centerline by skeletonization – in this case, a simple slice-wise centroid estimate from the segmentation.

Given such a centerline, we employed the following MATLAB-based, in-house segmentation method on the original, high resolution volumes:

1. Sub-volume extraction: Radial image resampling centered from each individual centerline point to construct a small fiber-unfolded sub-volume. Here, the axon-myelin interface can be seen as a surface with a strong negative gradient (going from bright axon to dark myelin).
2. Sub-volume layered segmentation: Perform layered surface segmentation on gradientbased cost-function of the sub-volume, using a graph cut approach^66^.
3. Volume labelling: Unfold the solution to the original volume coordinate system to obtain the final geometrically enforced solution.

The method is robust towards noise and image intensity variations. Furthermore, it is computationally efficient because the centerline serves to provide a much smaller sub-volume, in which we can build and solve the graph.

#### Segmentation of g-ratios along single internodes

Within the four overlapping XNH volumes depicted in Figure 2, we performed a rough manual segmentation of 6 axons with two visible nodes of Ranvier using the adaptive paintbrush in ITK-Snap, and extracted their centerlines by estimating their slice-wise centroids.

Thereafter, we generated *N* random points between the two nodes of Ranvier in MATLAB along each axon. We then extracted the relevant slices in which the randomly generated points could be found, and performed a manual segmentation of the inner axonal and outer myelin boundaries with the “roipoly” tool in MATLAB. The g-ratio estimation could be performed slice-wise and not perpendicular to the local axon trajectory because – as the ratio between the inner and outer axon boundaries – it is insensitive to skewed axons, unlike the equivalent AD.

### 4.4 Quantification of axonal dispersion of segmented axons

All calculations of axonal dispersion were based on the axonal centerlines. The point-to-point vectors within the centerlines of each of the 54 axons were averaged to produce a *main axon direction*. The average of the main axon directions across all 54 axons was defined as the *main bundle direction*. The OD of each axon is defined as the inclination angle between the main bundle direction and the main axon direction.

The axonal trajectories were significantly non-linear. To characterize how much the axons stray from their linear main directions, we introduced the concept of *micro-dispersion* at different length scales, *L*. To calculate the micro-dispersion, we first aligned the axons with the z-axis. Thereafter, the cumulative distance along the centerline was calculated, and the centerline was divided into segments of length, *L*, corresponding to the *sampling length*. A principal component analysis was performed on the set of center points within each segment to determine the *segment direction* in MATLAB. Thereafter, the inclination angle between the *segment direction* and the *main axon direction* was calculated. The average inclination angle across the segments (weighted according to the number of points in each segment) was defined to be the *mean inclination angle* of the axon at a length scale *L*. For a robust quantification, the axon skeleton was queried four times for each length scale *L* by shifting the starting position of the quantification by *L/4* as shown in Figure 4b. We studied length scales, *L*, ranging between 1 and 30 μm.

### 4.5 Monte Carlo simulations of the diffusion process in synthetic axons

Substrates G1-G5 in Figure 7 were based on the 54 segmented axons (substrate G6). Depending on their morphological properties, the 54 axons in geometries G1-G5 directly inherited one or more of the following from the segmented axons in G6: axon mean diameter, longitudinal diameter variation, orientation dispersion, micro-dispersion (trajectory). This was attained through modelling of the axons as deformed cylinders.

Prior to simulations, the reconstructed synchrotron meshes were post-processed to make them suitable for Monte-Carlo simulations of diffusing particles as follows. Firstly, the raw binary reconstructions were meshed using an octree-based surface extraction^67^ to ensure a closed surface with a high-polygonal density. Secondly, to reduce small irregularities arising from the surface reconstruction, the meshes were smoothed using an algorithm as in Desbrun et al. (1999)^68^, which ensures volume preservation while reducing high-frequency changes along the volumetric contour. Finally, the resulting meshes were triangulated and decimated to reduce the computational burden for the simulation.

All simulations were performed using the Monte Carlo Diffusion and Collision simulator presented in Rafael-Patino et al. (2020)^51^. Each axon in each geometry was simulated separately, and the intra-axonal diffusivity was set to 6.0 ·10^−10^ m^2^ s^−1^ as measured in Dyrby et al. (2013), as is conventional for ex vivo diffusion MRI. Particles were initialized uniformly within the central region of the axon meshes at a minimum distance of 20 μm from the axon ends. This confined initialization ensured that virtually no particle was able to diffuse outside the mesh. Perfectly elastic mesh boundaries were implemented, as in^51^. The number of particles and time-step duration (number of steps) were chosen using a bootstrap-based analysis of the convergence of the simulation as explained in^51^. In this investigation, we studied diffusion times, *D_t*, between 1-50 ms at intervals of 1 ms. This meant using 2·10^5^ particles and 5·10^5^ time steps with a duration of 1·10^−5^ s. The simulator output was in terms of the mean-squared-displacement, <*x*^2^>, of the particles in the directions parallel to and transverse to the main bundle direction.

The reported ADCs were calculated from <*x*^2^> by the Einstein relationship^69^:

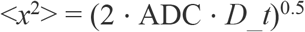

The AD estimations were based on the analytical expression for diffusion perpendicular to cylinders^70^: *λ*^2^ = *R*^2^/2 where *λ*^2^ is the mean-squared-displacement perpendicular to the cylinder and *R* is its radius.

