## Supplementary Figures 1 and 2 for "Axon morphology is modulated by the local environment and impacts the non-invasive investigation of its structure-function relationship"

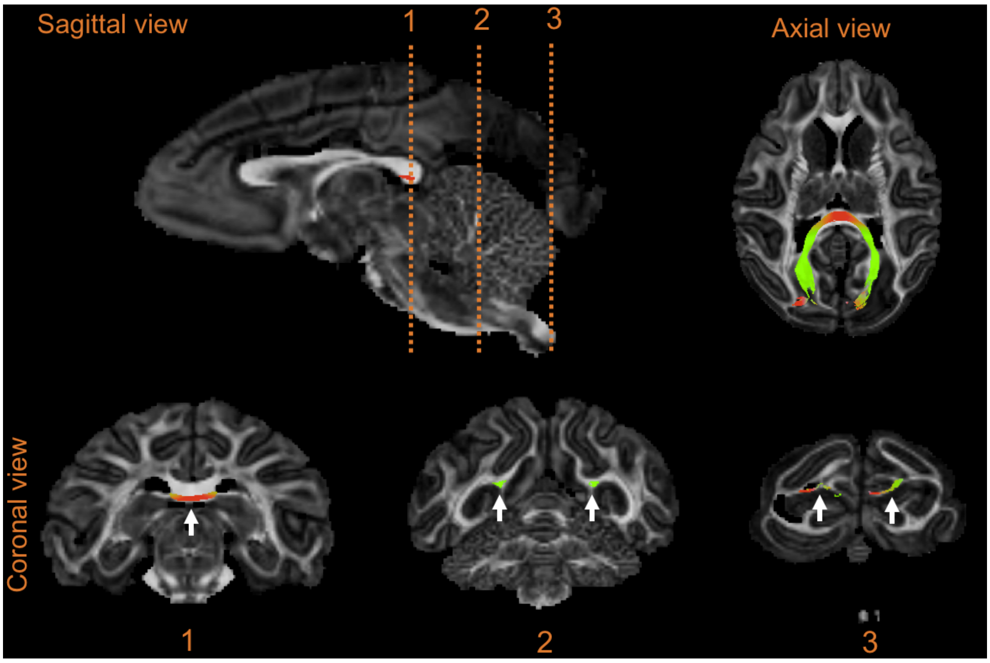


**Supplementary Figure 1.** Sagittal, axial and coronal views of monkey brain MRI volume and tractography of the interhemispheric connection between the V1/V2 visual cortices, passing through the splenium of the corpus callosum.

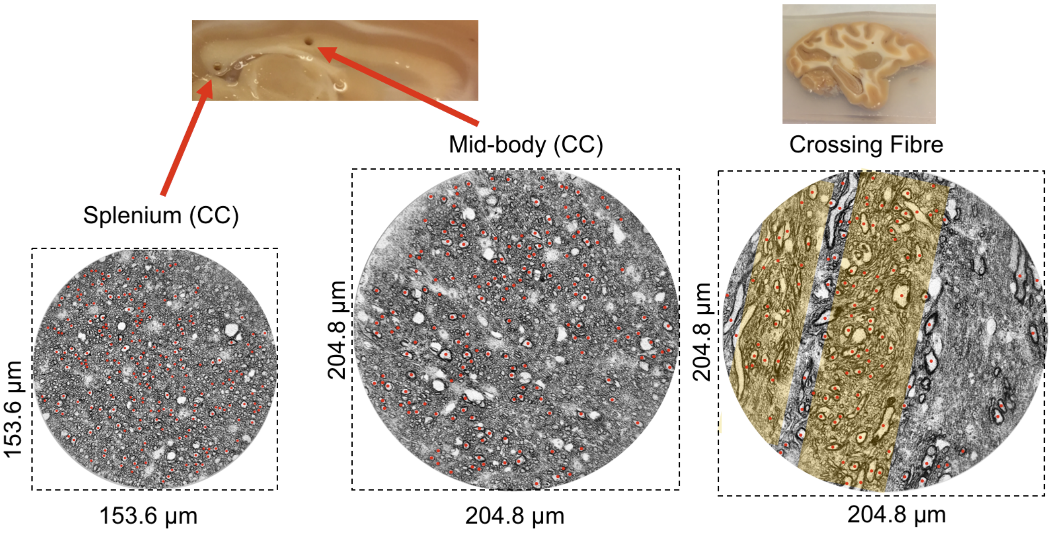


| | | | |
|---|---|---|---|
| Number density (large axons per $\mu m^2$) | 0.0123 | 0.0055 | 0.0038 |

**Supplementary Figure 2.** 2D cross sections of XNH-volumes from the splenium (75 nm voxel size), mid-body (100 nm voxel size) and crossing fiber (100 nm voxel size) regions. Axons with estimated equivalent diameters larger than 2 µm are marked by red dots. The large axons appear to be evenly distributed throughout the splenium and mid-body samples, but are organized in bands (yellow) in the crossing fiber region. Of the three samples, the number density of large axons is highest in the splenium and lowest in the mid-body. Photograph inserts show the positions at which 1 mm biopsies were extracted from the sagittal slices.
